# A pathogen-induced putative NAC transcription factor mediates leaf rust resistance in barley

**DOI:** 10.1101/2022.11.30.518475

**Authors:** Chunhong Chen, Matthias Jost, Megan A. Outram, Dorian Friendship, Sambasivam Periyannan, Jan Bartoš, Kateřina Holušová, Peng Zhang, Dhara Bhatt, Davinder Singh, Evans Lagudah, Robert Park, Peter Dracatos

## Abstract

Leaf rust, caused by *Puccinia hordei*, is one of the most widespread and damaging foliar diseases affecting barley (*Hordeum* spp.). The barley leaf rust resistance locus *Rph7*, located on the short arm of chromosome 3H, confers defence at all growth stages and was previously shown to have unusually high sequence and haplotype divergence. Earlier, four candidate genes for *Rph7* were reported and, despite an in-depth comparative sequence analysis and haplotypic characterisation, the causal gene could not be resolved. Here, we successfully cloned *Rph7* utilising a fine mapping approach in combination with an RNA-Seq based expression analysis. We identified three up-regulated and pathogen-induced genes with presence/absence variation (PAV) at this locus. Sequence analysis of chemically induced *Rph7* knockout mutant lines identified multiple independent non-synonymous variants, including a premature stop codon in a single non-canonical resistance gene that encodes a 302-amino acid protein. Progeny from four independent transgenic lines segregated for the expected avirulent *Rph7* infection type in response to several avirulent *P. hordei* pathotypes, however, all plants were susceptible to a single virulent pathotype confirming the specificity. Structural analysis using an AlphaFold2 protein model suggests that *Rph7* encodes a putative NAC transcription factor, as it shares structural similarity to ANAC019 from *Arabidopsis*, with a C-terminal BED domain. A global gene expression analysis suggests *Rph7* is involved in the activation of basal defence.

## Introduction

Cultivated barley (*Hordeum vulgare* L.) is the world’s fourth most important cereal, used primarily in malt production for alcoholic beverages and as grain feed for livestock and human food (Harwood, 2019). Of concern though are foliar diseases that reduce yield, grain quality and profitability (Cotterill et al., 1992). Leaf rust, caused by *Puccinia hordei* Otth., is widespread, resulting in regular seasonal epidemics and economically significant losses in global barley production (Park et al., 2015). Resistance to leaf rust in *Hordeum* spp. is well characterised and widely available due to the numerous genetic (Dracatos et al., 2015; Elmansour et al., 2017; Jost et al., 2020; Mehnaz et al., 2021; Mehnaz et al., 2022; Park *et al*., 2015; Qi et al., 1998; Rothwell et al., 2020; Singh et al., 2016; Singh et al., 2018) and cloning studies undertaken (Chen et al., 2020; Dinh et al., 2022; Dracatos et al., 2019; Wang et al., 2019). However, in most cases the underlying genes responsible for resistance are not known, limiting effective and efficient deployment in agricultural settings (Dinh et al., 2020; Park et al., 2015).

The plant immune system is multi-layered, with both extra- and intracellular receptors being critical for pathogen recognition and disease resistance signalling (Zhou and Zhang, 2020). Race-specific resistance, referring to defence against certain races of a pathogen, occurs via perception of the pathogen governed by two predominant immune receptor families: nucleotide binding-leucine-rich repeat (NLRs) and receptor-like kinase (RLKs) receptors. To date, the most prevalent immune receptors characterised in cereal crops are NLRs. However, recent evidence suggests race-specific resistance is also governed by non-canonical gene classes (Dinh et al., 2020; Sánchez-Martín and Keller, 2021; Zhang et al., 2020). Two recent studies utilised sequenced chromosome scale assemblies from the wheat and barley pangenomes to clone the race-specific leaf rust resistance genes *Lr14a* (Kolodziej et al., 2021) and *Rph3* (Dinh et al., 2022). Interestingly, *Lr14a* and *Rph3* encode a membrane-bound ankyrin repeat containing protein and a putative executor protein, respectively. These studies highlight opportunities to both explore and exploit diverse resistance mechanisms in cereal crops.

*Rph7* is a semi-dominant inherited gene, first described from the cultivar ‘Cebada Capa’ (PI 539113) (Roane and Starling, 1970). *Rph7* confers all-stage resistance against most *P. hordei* isolates in Europe and Australia, though isolates virulent on *Rph7* have been identified in Spain (Shtaya et al., 2006), the Near East (Golan et al., 1978), North America (Steffenson, 1993), and most recently in Australia. Trisomic analysis initially mapped *Rph7* to barley chromosome 3H (Tuleen and McDaniel, 1971) and subsequent biparental mapping further refined the *Rph7* locus to the short arm of chromosome 3H, near the telomere (Graner et al., 2000). Fine mapping and sequencing of physically overlapping BAC clones spanning the determined interval revealed that *Rph7* is located within a chromosomal region of high haplotypic divergence, largely explained by the presence of numerous cultivar-specific insertions containing different classes of retrotransposons (Brunner et al., 2003; Scherrer et al., 2005). Shotgun Sanger sequencing and assembling of BAC clones of the leaf rust susceptible cultivar Morex and the *Rph7*-containing line Cebada Capa revealed the presence of a 100 kb sequence insertion in Cebada Capa while the genes at the *Rph7* locus were conserved (Scherrer et al., 2005). No typical resistance gene candidates were identified within the cultivar-specific insertion. Therefore, four co-segregating genes (*HvHGA1, HvHGA2, HvPG1* and *HvPG4*) were considered as the most logical candidates for the *Rph7* resistance. Despite an in-depth comparative sequence analysis and haplotypic characterisation at the *Rph7* locus, the causal gene underpinning the resistance was not resolved.

Here, we built on the evidence of Scherrer *et al*. (2005) with the aim of determining the molecular basis of *Rph7* resistance in Cebada Capa. We performed further fine mapping to develop additional recombinants at the *Rph7* locus to either confirm or rule out the four previously determined candidates. We subsequently used RNA-Seq to further examine differential gene expression at two different time points following infection using an Australian pathotype of *P. hordei* avirulent for *Rph7*. Confirmation of several pathogen-induced gene candidates was then performed using chemical mutagenesis and complementation experiments based on detailed rust testing with *P. hordei* pathotypes with differential infection responses on *Rph7*-carrying lines. Finally, we used structural prediction and analysis, as well as a global gene expression analysis to gain insights into the putative function of the identified resistance gene.

## Results and Discussion

Recent crop pangenome projects have revealed both the extent and importance of intraspecies polymorphisms, including presence-absence variations (PAVs), highlighting the inadequacy of previous over-reliance on reference genome information (Walkowiak et al., 2020). PAVs are especially relevant for resistance gene classes that evolve via duplication and diversifying selection (like NLRs). Often the causal resistance gene is either absent or partially represented in susceptible accessions (Sánchez-Martín et al., 2016). To resolve the underlying molecular basis of the *Rph7* resistance we tested two hypotheses. The first was that the involvement of one or more of the previously postulated candidate genes (*HvPG1, HvPG4, HvHGA1* and *HvHGA2)* could be confirmed or eliminated through additional recombinant screening at the *Rph7* locus. The second was that an RNA-Seq based expression analysis would reveal additional candidate genes at the *Rph7* locus. We therefore performed additional genetic fine mapping, using an alternative susceptible barley genotype Wabar2722 (*rph7*) crossed with the *Rph7* donor line Cebada Capa (*Rph7*), to test whether we could identify further recombinants in the previously defined genetic interval. Newly identified recombinants eliminated the involvement of three of the previous *Rph7* gene candidates (*HvHGA1, HvHGA2* and *HvPG4*) (Figure 1A, Supplemental Table 1).

**Figure 1:**
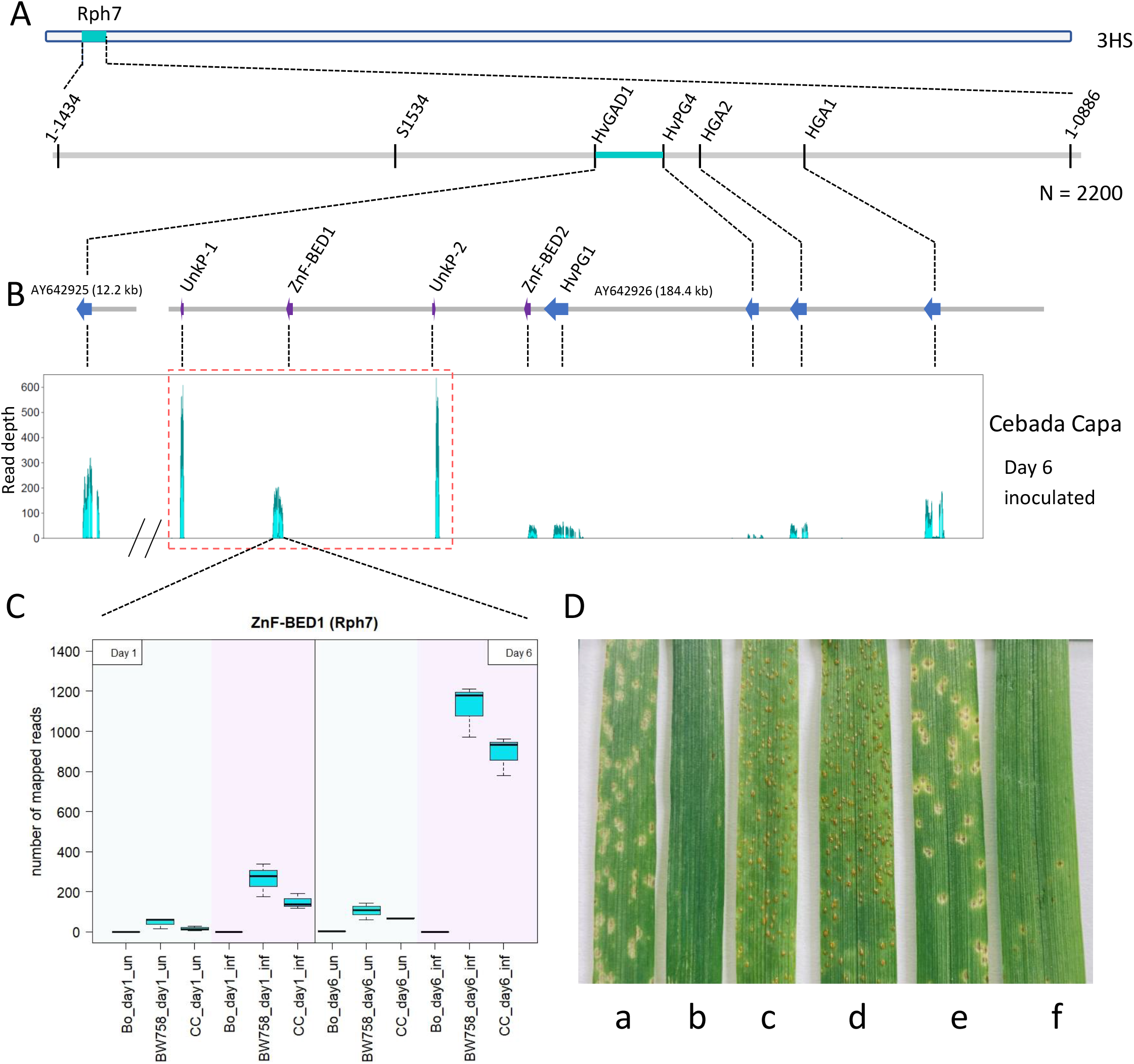
Gene discovery at the *Rph7* locus In Cebada Capa. A) Genetic mapping narrowed the *Rph7* locus between flanking markers within genes *HvGAD1* and *HvPG4* B) Schematical overview of previous sequenced BAC contigs (grey, Scherrer et al. 2005). RNA-Seq data revealed the gene expression of previous reported candidate genes (blue) and identified further four expressed candidate genes labelled in purple. Among the four newly identified expressed candidates, three genes are upregulated after infection (highlighted in red box) in the resistant lines Cebada Capa and the near isogenic line BW758 (Bowman+*Rph7)* compared to susceptible Bowman at day 6. (C) Expression profile of *Znf-BED1* (*Rph7*). Samples labelled with light blue background were treated as mock control, while purple background labelled shows the expression patterns are from barley leaves infected with *Rph7*-avirulent *Puccinia hordei* pathotype 5457 P+ for Bo (Bowman), BW758 and CC (Cebada Capa). D) Complementation results showing phenotypic responses at the seedling stage 10 days after infection with *Puccinia hordei* pathotypes 5477 P-(a, c, d, and e) and 200 P+ (b and f) that elicit intermediate and low infection types respectively. From L to R (a and b) resistant sib of the T1 transgenic Golden Promise + *Rph7* line B114-1, (c) leaf rust susceptible controls Golden Promise and (d) Bowman and (e and f) *Rph7* carrying positive control BW758.

We performed an RNA-Seq experiment at the seedling stage for an early (24 hours) and late (day 6) timepoints, to determine the genes specifically expressed during infection when challenged with an *Rph7* avirulent *P. hordei* isolate. We identified five expressed genes within the target interval, four of which were not predicted in the previous *Rph7* study (Scherrer *et al*., 2005). Interestingly, three of the genes (viz. *UnkP-1, ZnF-BED1* and *UnkP-2*) were upregulated only in the *Rph7* carrying lines Cebada Capa and BW758 (near isogenic line carrying *Rph7* in the susceptible cv Bowman background) (Figure 1B and 1C, Supplemental Table 2). This suggests one or more of these genes may mediate *Rph7* resistance in Cebada Capa. To further verify the involvement of the expressed candidate genes at the locus we chemically mutagenized the BW758 line and progeny-tested eight susceptible knockout M_4_ families.Sanger sequencing of the five expressed genes within the target *Rph7* interval (*UnkP-1, ZnF-BED-1, UnkP-2, ZnF-BED-2* and *HvPG1*) on all progeny-tested susceptible mutants determined that four out of eight mutant lines contained chemically induced SNPs within the coding region (either C>T or G>A) of the *ZnF-BED-1* gene (Supplemental Figure 1A). No non-synonymous mutations were identified in the remaining four mutant lines in *ZnF-BED1* or in any of the sequenced neighbouring genes. To determine whether further candidates could be identified carrying SNPs in the missing genes of the sequence gap between the two BAC clones we sorted and sequenced chromosome 3H from the wild type and mutants. MutChromSeq analysis confirmed *ZnF-BED1* as the primary candidate and confirmed no additional plausible candidates on chromosome 3H. This suggests the possible presence of mutations in downstream targets of *Rph7* or genes regulating the expression of resistance.

The genomic sequence structure of *ZnF-BED1* consists of 1,197 nucleotides, including four exons and three introns, which encode for a 302-residue protein with a sequence-predicted zinc-finger BED domain (ZnF-BED) at the C-terminus (Figure 2C). To conclusively confirm the involvement of *ZnF-BED1* in mediating *Rph7* resistance we performed a complementation experiment by cloning a 3,625pb genomic fragment including the native promoter and terminator into Golden Promise using *Agrobacterium*-mediated transformation. Rust testing was performed on T_1_ generation Golden Promise + *ZnF-BED1* lines and controls Golden Promise, Bowman, BW758 and Cebada Capa using four *P. hordei* pathotypes eliciting differential responses on *Rph7* carrying lines (Supplemental Tables 3 and 4). Four T_1_ lines segregated for a single gene (3 Resistant:1 Susceptible) for the expected *Rph7* phenotype in response to three *Rph7* avirulent pathotypes. In contrast, the same four T_1_ lines were susceptible to the *Rph7*-virulent pathotype as the resistant controls confirming the *Rph7* specificity of the Golden Promise transgenic lines. Further genotypic analysis using PCR markers designed to the selectable marker and *ZnF-BED1* was performed to confirm the transgenic status of the lines (Supplemental Table 4). The correlation of segregation patterns observed in the T_1_ lines (B114-1, B114-2, B114-17, and B114-18) in response to all *P. hordei* pathotypes suggests that a single resistance factor (*ZnF-BED1)* is sufficient to confer *Rph7* mediated resistance. We therefore refer to *ZnF-BED1* as *Rph7* for the remainder of the manuscript.

**Figure 2:**
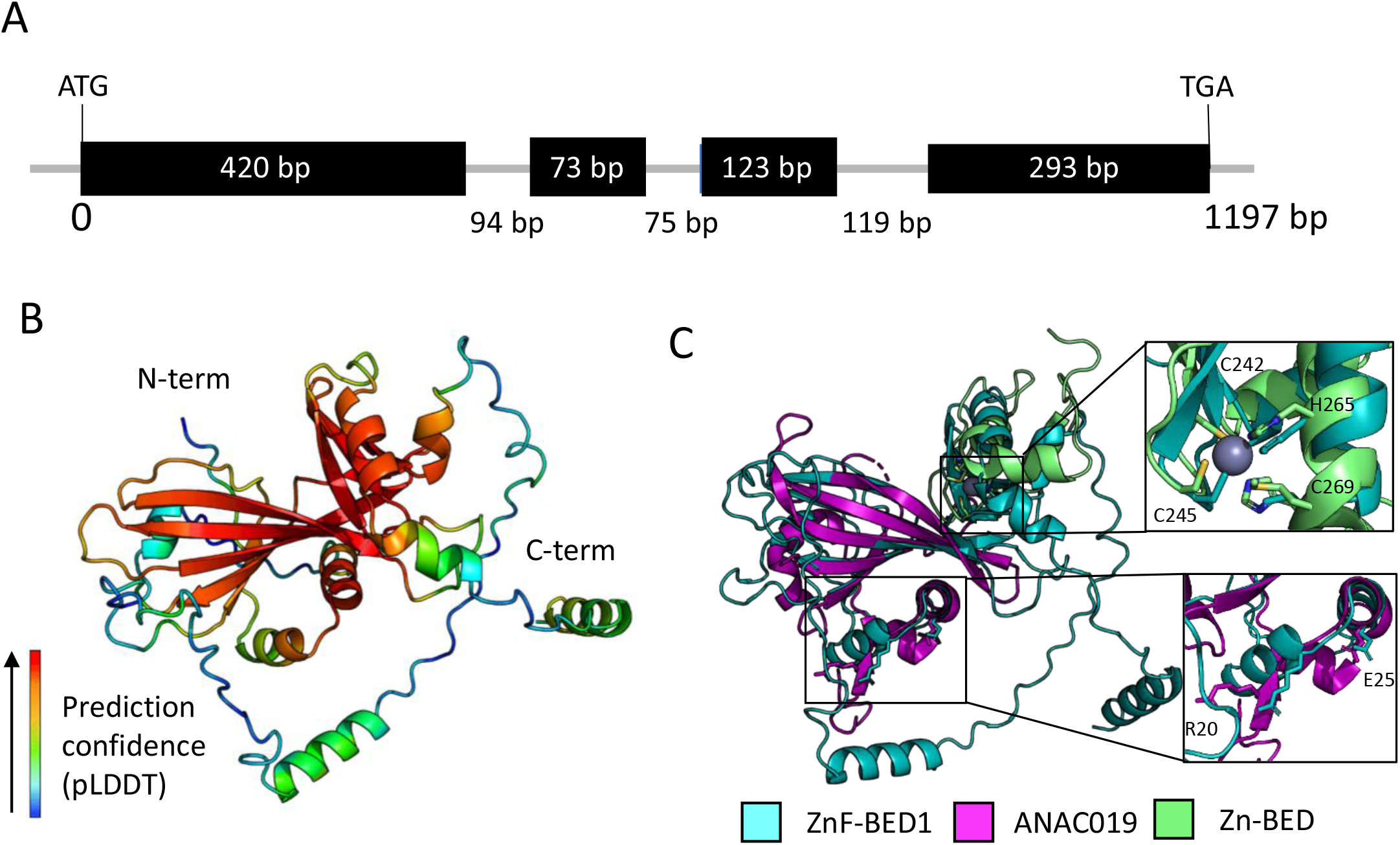
ZnF-BED1 shares structural similarity with plant specific NAC domain-containing proteins. (A) Schematical drawing of gene structure of Znf-BED1 consisting of 4 exons (black boxes) and 3 introns (grey) with respective length in bp. (B) AlphaFold2 prediction of ZnF-BED1 shown as cartoon representation and coloured by per-residue confidence score (pLDDT), where red is the most confident and blue is the least confident. (C) Structural superimposition of the AlphaFold2 prediction of ZnF-BED1 (teal) and top structural match from Dali ANAC019 NAC domain (PDB ID: 3SWP; magenta) shows similarity is limited to the N-terminus of ZnF-BED, and superimposition of the top structural match from Dali for the C-terminal domain (residues 220-302), the C2H2 type zinc finger domain of human zinc finger BED domain containing protein 2 (PDB ID: 2DJR, green). Insert shows residues involved in Zn co-ordination in 2DJR, and dimerization in 3SWP, and corresponding putative residues that may be involved in ZnF-BED1. Residues are labelled according to ZnF-BED1.

Sequence comparison across the five expressed genes at the locus between Cebada Capa and a further five barley cultivars predicted to carry *Rph7* (Ellinor, Toddy, Galaxy, Dictator 2 and La Estanzuela), confirmed they all shared the same haplotype. Further haplotypic comparisons between Cebada Capa and the 20 sequenced accessions comprising the barley pangenome (Jayakodi et al., 2020) identified three distinct haplotypic groups: H1-H3. each varying in gene content corroborating previous haplotypic data reported by Scherrer *et al*. (2005). H1 was similar structurally to Cebada Capa, H2 contained mostly truncated homologs of only *Rph7* and *UnkP-1*, whereas H3 accessions lacked all pathogen induced genes at the *Rph7* locus (Supplemental Figure 2). Accessions Barke and Hockett shared the same full-length transcribed protein sequence, whereas RGT Planet and HOR3365 contain a splice site variant and the remaining accessions had variable partial variants of *Rph7*. Further investigation of the genomic sequence between Cebada Capa and the four accessions with full length homologues of the *Rph7* gene revealed numerous SNP and indel polymorphisms in the predicted putative NAC domain at the N-terminus and a splice site mutation in the intron-exon boundary. These comparisons suggest that resistance is likely due to either a PAV of the *Rph7* gene or the presence of a splice site mutation at the end of exon one leading to a truncated protein in those accessions that carry a homologue (Supplementary Figure 3).

To investigate the molecular function of *Rph7* we predicted the protein structure using AlphaFold2 (Jumper et al., 2021). This showed that Rph7 consists predominantly of a central β-sheet surrounded by α-helices (Figure 2B). Using the top ranked model (Figure 2B), which has an average pLDDT (predicted lDDT-Cα; per-residue measure of local confidence) score of ∼70, we performed a structural search against the protein databank (PDB) using the Dali server (Holm, 2022). Dali reports structural similarity by Z-score, where significant similarities are indicated by Z-scores >2. Four of the top five unique proteins (Z-score > 6) were NAC (NAM, ATAF1,2, and CUC2) domain-containing proteins (Supplemental Table 5). Rph7 and the DNA-binding NAC domain of *Arabidopsis* ANAC019 (the top structural match) share ∼20% sequence identity (Supplemental Table 5). Structural superimposition between the two proteins shows the structural similarity to ANAC019 is limited to the N-terminus of Rph7 (Figure 2C). To confirm our previous observation that the C-terminus is a putative zinc-finger BED-domain, we did a structural search with the C-terminal region alone (residues 220-302). Unsurprisingly we found the top structural hits were C2H2-type ZnF domains, and it appears that a zinc co-ordination motif is present in Rph7 (Figure 2C). Taken together, Rph7 contains an N-terminal NAC domain and C-terminal zinc-finger BED domain separated by a long-disordered loop. NAC proteins typically consist of a conserved ∼150 amino acid N-terminal NAC domain that is capable of binding DNA and facilitates dimerization, and a diverse C-terminal domain that typically functions as a transcription regulatory domain (Welner et al., 2012). In *Arabidopsis*, ANAC019 has a largely positively charged surface patch due to a cluster of arginine and predominantly lysine residues that are responsible for interacting with the backbone phosphates of the DNA molecule. Similarly, Rph7 shows a positive surface suggesting that like ANAC019 it may be capable of binding DNA (Supplemental Figure 4) (Welner *et al*., 2012). Dimerization in ANAC019 is largely mediated by two prominent salt bridges formed by conserved R19 and E26 resides, which localise in a similar region on the predicted Rph7 model (Figure 2C). As such, it is tempting to speculate that the NAC domain of Rph7 dimerises and binds to DNA in the same manner as ANAC019 though this remains to be experimentally determined.

To further understand the implications of the previously identified chemically induced mutants we mapped each of the sequence confirmed Rph7 mutants (G72E, T90I, R188*, and D209N) to the predicted structure (Supplemental Figure 1B). All four Rph7 mutants were surface exposed in the model and localise predominantly to flexible regions within the protein (Supplemental Figure 1B). The introduction of a premature stop codon at R188 would result in the production only of the NAC-domain with an extended C-terminus, suggesting that it would retain the capacity to bind to DNA and presumably oligomerise but lacks the C-terminal ZnF-BED/regulatory domain. D209 occurs within the flexible linker between the NAC and BED domain. Two of the mutants G72E and T90I localise to the opposite side of the protein, away from the putative DNA-binding surface suggesting these mutants would likely not impact this function directly. Though G72 localises close to R20 and E25 in the structure, it perhaps could be involved in mediating dimerization, should Rph7 function similarly to ANAC019.

Plant basal defence involves pathogenesis-related (*PR*) gene expression mediated by transcription factors (TFs). In *Arabidopsis* stress-responsive NAC TFs respond to phytohormones, such as salicylic and jasmonic acid at the infection site, resulting in the expression of *PR* genes to induce the production of antifungal proteins and enzymes. We used the RNA-Seq data generated in this study to perform a differential gene expression (DEG) analysis comparing the transcriptomes of Bowman and the BW758 near isogenic line (NIL) from the two timepoints. Unsurprisingly DEGs were substantially higher at day 6 in BW758 mirroring the expression of *ZnF-BED1* in *Rph7*-carrying lines (Figure 3A). In parallel, Gene Ontology enrichment analysis revealed multiple biological processes related to the activation of basal plant defence (Supplemental Figure 5). We focussed on four main classes of DEGs specific to fungal attack identified in BW758, including Jasmonate-related (*JR*), pathogenesis related (*PR*), *WRKY* transcription factors and *NLR* genes (Supplemental Figure 6). In all cases *JR* and *PR* genes were up regulated in response to infection, suggesting, as previously reported, that NAC TFs like Rph7 play an important role in regulating or modulating the cellular plant defence responses.

**Figure 3:**
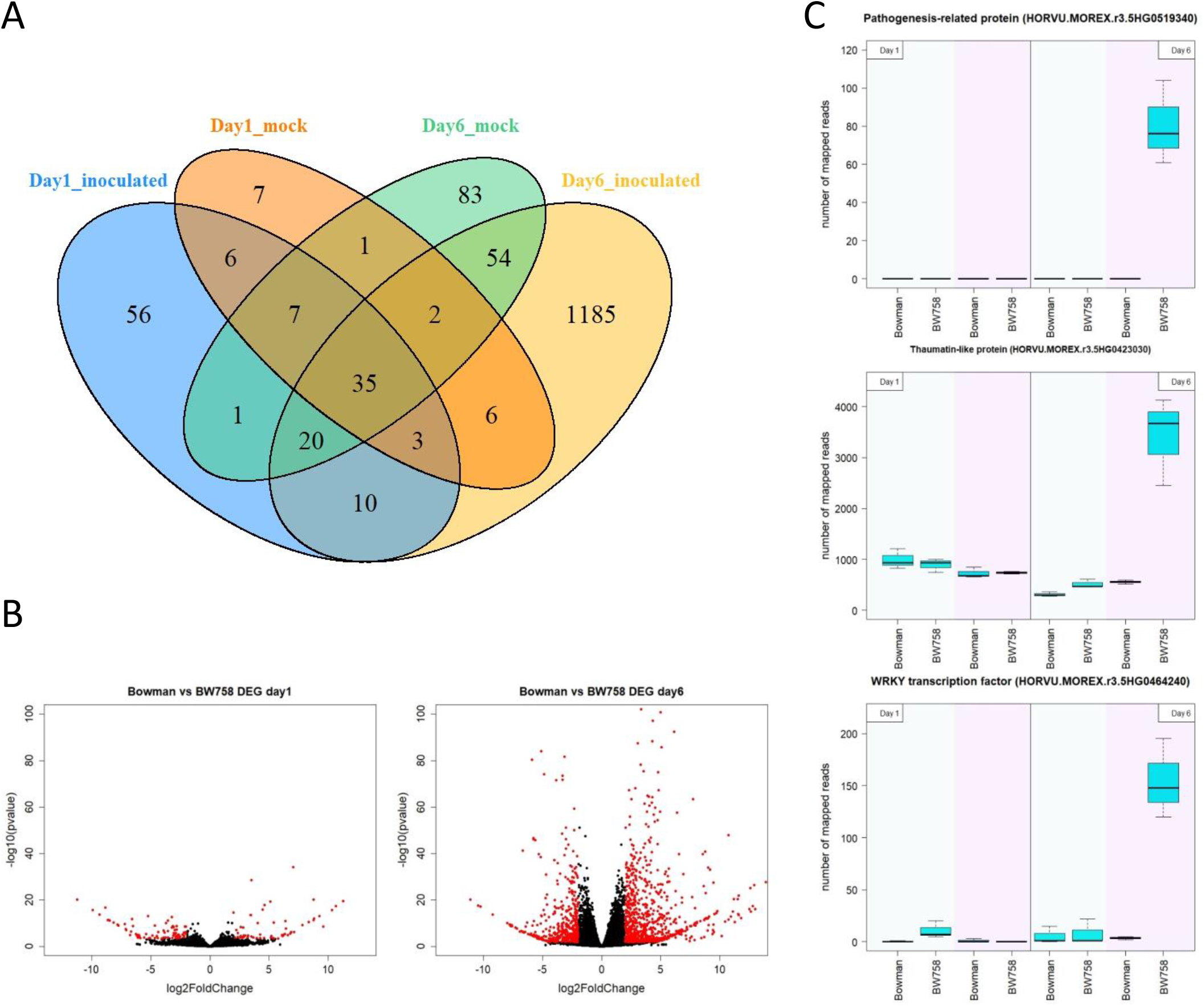
Differential Expressed Genes (DEGs) detected between Bowman and BW758 (Bowman+ *Rph7*). A) Overview of the number of DEGs detected for treated (infected with *Rph7* avirulent *Puccinia hordei* pathotype 5457 P+) and untreated (mock oil) at day 1 and day 6 after inoculation. B) Volcano plots showing detected DEGs at day 1 (left) and day 6 (right). DEGs with Log2Fold change < or > 2 are highlighted in red. C) Examples of detected key disease response marker genes showing upregulation at day 6 of infection that are predicted to encode pathogenesis related protein, thaumatin-like protein and a WRKY transcription factor. A full list of DEGs involved in basal plant defence were provided in supplemental Figure 6.

Despite the characteristic near immune response elicited by most *Rph7* avirulent *P. hordei* pathotypes, surveillance studies in Australia identified a group of pathotypes that elicited a necrotic intermediate response, and more recently, a single isolate that was fully virulent on *Rph7*. This led to the hypotheses that either two or more genes may confer *Rph7*-mediated resistance or alternatively that distinct *P. hordei* isolates eliciting a differential *Rph7* response were either homozygous or heterozygous for the avirulence gene matching *Rph7*. Our data rules out the two gene hypothesis, however, genetic analyses based on crosses of *P. hordei* pathotypes with contrast (immune x intermediate *Rph7* response) and comparative sequence analysis of the corresponding *Rph7* cognate effector are required to test the heterozygous vs homozygous avirulence hypothesis. Further mechanistic studies on the downstream targets and regulation of the *Rph7* resistance will best equip biotechnologists to efficiently engineer crop plants for durable disease control.

## Methods

### Plant material

High resolution F_2_ fine mapping population (n=2200 gametes) was developed by intercrossing Cebada Capa (*Rph7*) with WABAR 2722 (*rph7*). The NIL BW758 (Bowman+*Rph7*) was used for mutagenesis as well differential gene expression experiments in comparison to the wild type cv Bowman.

### Pathogen materials and phenotypic analysis

Details of the four *P. hordei* pathotypes used in this study, including pathogenicity on different resistance genes, are listed in Supplemental Table 3. Pathotypes were designated according to the octal notation proposed by Gilmour (1973) and Park and Karakousis (2002). Pathotype 5457 P+ was used to phenotype the mapping population for recombinants, screening mutants and RNA-Seq expression analysis. *Rph7*-avirulent (200 P+, 276 P+ and 5477 P-) and virulent (5553 P+) pathotypes were used to validate the resistance gene function and specificity in the T_1_ generation of Golden Promise + *Rph7* transgenic plants. These pathotypes were isolated and increased as described by Dinh *et al*. (2022) before being stored in liquid nitrogen at the Plant Breeding Institute, the University of Sydney, Australia. Rust testing of barley seedlings in the greenhouse with *P. hordei* pathotypes listed above was performed as described in Dinh *et al*. (2022) and plants were phenotyped 10 days post-inoculation using the “0”– “4” infection type (IT) scale as described by (Park et al., 2015).

### Mutagenesis

A mutant population was developed by chemically treating 1,000 BW758 seeds using sodium azide as described by Chandler and Harding (2013) with some modifications. Seeds were wrapped in cheesecloth, immersed in water at 4°C overnight and then transferred into 2-litres of water to aerate with pressurised air for 8 hr followed by draining. Seeds were treated on a shaker with 1 mM sodium azide dissolved in 0.1 M sodium citrate buffer (pH 3.0) on a shaker for 2 hr, washed under running water for at least 2 hr, and then dried in a fume hood overnight. Seeds were space planted in the field directly and harvested as described by Chen et al. (2021). More than 4,500 M_2_ generation spikes were phenotyped for segregating *rph7* knockouts, as described previously (Chen *et al*., 2021; Dracatos *et al*., 2019). Ten M_2_ families were identified as putative *Rph7* knockout mutants. Subsequent progeny testing of M_3_ families confirmed seven homozygous susceptible and one segregating family for further sequence analysis. For MutChromSeq analysis, non-amplified DNA of 3H chromosome of BW758 and seven susceptible mutants was shotgun sequenced after flow-cytometric sorting as described previously (Chen et al., 2021; Dracatos et al., 2019). Wild-type reference sequence was assembled using Meraculous (Chapman et al., 2011) and candidate gene identification was done according to Sánchez-Martín *et al*. (2016). All raw reads produced were submitted to SRA archive under BioProject ID PRJNA906712.

### RNA-Seq data preparation

Cebada Capa, Bowman + *Rph7* (BW758) and Bowman seedlings were grown in trays as described by Dinh *et al*. (2022) and then inoculated using either oil (mock control) and oil mixed with 30 mg of *P. hordei* urediniospores (pt. 5457 P+). Leaf tissue (three technical replicates for each of the three biological replicates per genotype) was harvested 24 hours (day 1) and 6 days after inoculation (dai) and snap frozen in liquid N and stored at -80°C (Supplemental Table 6). RNA was isolated using the Maxwell robot (Promega). RNA quality and quantity was assessed by RNA gel electrophoresis and the Nanodrop spectrophotometer. Illumina paired end 150 bp sequencing was performed by Novogene yielding between 21 and 37 million raw reads per sample. Adapter and quality trimming of raw reads was performed using fastqc version 0.11.9 (Andrews, 2010) and trimmomatic 0.39 (Bolger et al., 2014), respectively.

### Candidate gene identification

Genetic screen for recombinants has been performed with PCR based markers or KASP markers (Figure 1A, Supplemental Table 7). Critical recombinants have been evaluated in F_3_ family phenotypically and genotypically (Supplemental Table 1). The sequence section between the conserved genes *HvGAD1* and *HvHGA1* of the Morex_V2 (Monat et al., 2019) reference sequence has been replaced with sequenced BACs AY642925 and AY642926 (Scherrer *et al*., 2005) to avoid background mapping of similar reads within the transcriptome sequences. Cleaned RNA-Seq reads were mapped with Tophat 2.1.1 (Kim et al., 2013) and Bowtie2 2.4.4 (Langmead and Salzberg, 2012) using default settings. Read depth for each base pair was calculated with the depth function of SAMtools version 1.12 (Danecek et al., 2021). Mean read coverage of biological replicates were visualized with R bar plot to identify expressed sequence regions.

### In silico expression analysis

*De novo* transcriptome assembly of Cebada Capa was performed with trinity version 2.13.2 (https://github.com/trinityrnaseq/trinityrnaseq, Grabherr et al. (2011)). We selected one replicate from day 1 and day 6 of infected and uninfected Cebada Capa to generate a Cebada Capa specific reference (B4, B27, C4 and C28, Supplemental Table 6). These were pooled before assembly. Gene and isoform expression levels were estimated with RSEM version 1.3.3 (Li and Dewey, 2011) using trinity mode (https://github.com/deweylab/RSEM/releases). Contigs of the trinity assembly have been matched with candidate genes by BLASTn and read counts visualized by R boxplot.

### Differential gene expression analysis

Purified RNA-Seq reads have been mapped against Morex_V3 (Mascher et al., 2021) with HiSat2 version 2.2.1 (Kim et al., 2019). BAM alignment files have been sorted with SAMtools 1.12 (https://github.com/samtools/samtools/, Danecek *et al*. (2021). Reads mapped to transcripts using gff3 file of Morex v3 annotation (July 2020, Mascher et al., 2021) by ht-seq-count (Anders et al., 2015). Differential gene expression analysis has been performed with DEseq2 R package (Love et al., 2014). Genes expressed with a log2 fold change >+/-2 have been subjected to gene ontology enrichment using the Triticeae-Gene Tribe database (http://wheat.cau.edu.cn/TGT/) (Chen et al., 2020).

### Candidate gene cloning, vector construction and barley transformation

Primers were designed to amplify the *ZnF-BED1* gene from Cebada Capa including the sequence of the native promoter and terminator that are 1.5kb upstream of the ATG and 1kb downstream of the STOP codon respectively. Cloning and *Agrobacterium*-mediated transformation of a 3,625 bp genomic fragment was performed as described in Chen *et al*. (2021).

### Structural modelling and structural comparisons of ZnF-BED1

Five structural models of ZnF-BED1 were generated using Google DeepMind’s AlphaFold2 (Jumper et al., 2021). Full databases were used for multiple sequence alignment (MSA) construction. All templates downloaded on July 20, 2021, were allowed for structural modelling. We selected the best model (ranked_0.pdb) for downstream protein visualisation using open-source Pymol or UCSF *ChimeraX* (Pettersen et al., 2021). Structural comparisons to known structures in the protein databank were carried out using the Dali webserver (Holm, 2022).

## Acknowledgements

The authors thank Beat Keller and Gerhard Herren for scientific discussion about details of the previous published work. We thank B. Clark, L. Ma, M. Williams, and S. Hoxha for technical supports with plant growth, molecular experiments, mutagenesis, and pathogen spore increases. We thank the GRDC for supporting the research, PD, CC, MJ, DF were funded under grant UOS1507-005RMX - US00074. PD was also supported by La Trobe University and the Alexander von Humboldt Foundation.

## Figure legends

**Supplemental Figure 1:** Chemically induced SNPs identified in the coding sequence of *ZnF-BED1*. A) Structure of the candidate *ZnF-BED1* gene from Cebada Capa showing the position of four independent sodium azide-induced mutants within the coding sequence indicating that the *ZnF-BED1* gene was required for *Rph7* mediated resistance. B) Transparent surface representation of ZnF-BED1 highlighting the four non-synonymous amino acid changes identified in ZnF-BED-1 mutant lines. Residues are shown in stick representation and coloured in orange. C) Phenotypic responses at the seedling stage 10 days after infection with *Rph7*-avirulent *Puccinia hordei* pathotype 5457 P+ of resistant WT (BW758) and four independent mutants carrying either G to A or C to T mutations in the *ZnF-BED1* gene.

**Supplemental Figure 2:** Pan-Genomic representation of expressed genes detected within the genetic interval between *HvGAD1* and *HvPG4* at the *Rph7* locus. Haplotype 1 (H1) shows the highest level of structural conservation to the *Rph7* resistance haplotype in Cebada Capa evidenced by size and gene content. H2 accessions carry a homolog of *Rph7* but are missing either one or more of the other expressed genes within the interval. H3 accessions have no homologous sequence to *Rph7*, or the other Cebada Capa expressed genes, which is evidenced by a reduced interval size indicative of a presence/absence variation.

**Supplemental Figure 3:** Sequence alignment of genomic sequence of the full-length *Rph7* homologs detected in the barley Pan Genome. The top row represents the full-length genomic sequence of the *Rph7* candidate gene *ZnF-BED1* in resistant donor cultivar Cebada Capa with exons highlighted in blue. Sequence variations are restricted to the N-terminal part of the gene within the predicted NAC domain. The homolog sequences carry a SNP mutation at the end of Exon 1 (highlighted in red) which likely causes an alternated splicing variant and does not translate to a full-length protein with similarity to *Rph7*. The remaining four partial homologs (in accessions HOR10350, HOR9043, Golden Promise and OUN333) cover the last 793 bp conserved with the displayed sequences and are not shown in this supplementary figure.

**Supplemental Figure 4:** ZnF-BED1 has a large positively charged surface patch suggesting it may bind DNA. Surface representation of AlphaFold prediction of ZnF-BED1 (left) and Arabidopsis ANAC019 (right, 3SWP) showing electrostatic charge generated with Chimera X. Blue indicates positive charge, red negative charge, and white is neutral.

**Supplemental Figure 5:** Gene ontology enrichment analysis of detected differential expressed genes (DEG) between Bowman (susceptible) vs BW758 (resistant) at day 1 (A) and day 6 (B) after inoculation with *Rph7*-avirulent *Puccinia hordei* pathotype 5457 P+. Overview of Biological Processes detected in DEGs using Triticeae Gene Tribe (http://wheat.cau.edu.cn/TGT/).

**Supplemental Figure 6:** Subset of day 6 differential expressed genes (DEGs) indicating *ZnF-BED1* (*Rph7*) is likely involved in activating the basal disease response. DEGs between Bowman and BW785 are colour coded according to log2FoldChange value for upregulated genes in red and downregulated genes in blue. Gene descriptions have been extracted from Morex_V3 (Jul 2020) annotation file.

## References

Anders, S., Pyl, P.T., and Huber, W. (2015). HTSeq—a Python framework to work with high-throughput sequencing data. Bioinformatics 31:166–169. 10.1093/bioinformatics/btu638.

Andrews, S. (2010). FastQC: A quality control tool for high throughput sequence data https://www.bioinformatics.babraham.ac.uk/projects/fastqc/.

Bolger, A.M., Lohse, M., and Usadel, B. (2014). Trimmomatic: a flexible trimmer for Illumina sequence data. Bioinformatics 30:2114–2120. 10.1093/bioinformatics/btu170.

Brunner, S., Keller, B., and Feuillet, C. (2003). A large rearrangement involving genes and low-copy DNA interrupts the microcollinearity between rice and barley at the Rph7 locus. Genetics 164:673–683. 10.1093/genetics/164.2.673.

Chandler, P.M., and Harding, C.A. (2013). ‘Overgrowth’ mutants in barley and wheat: new alleles and phenotypes of the ‘Green Revolution’ DELLA gene. Journal of experimental botany 64:1603–1613. 10.1093/jxb/ert022.

Chen, C., Jost, M., Clark, B., Martin, M., Matny, O., Steffenson, B.J., Franckowiak, J.D., Mascher, M., Singh, D., Perovic, D., et al. (2021). BED domain-containing NLR from wild barley confers resistance to leaf rust. Plant Biotechnology Journal 19:1206–1215. 10.1111/pbi.13542.

Chen, Y., Song, W., Xie, X., Wang, Z., Guan, P., Peng, H., Jiao, Y., Ni, Z., Sun, Q., and Guo, W. (2020). A collinearity-incorporating homology inference dtrategy for vonnecting emerging assemblies in the triticeae tribe as a pilot practice in the plant pangenomic era. Molecular Plant 13:1694–1708. 10.1016/j.molp.2020.09.019.

Cotterill, P., Rees, R., Platz, G., and Dill-Macky, R. (1992). Effects of leaf rust on selected Australian barleys. Australian Journal of Experimental Agriculture 32:747–751. 10.1071/EA9920747.

Danecek, P., Bonfield, J.K., Liddle, J., Marshall, J., Ohan, V., Pollard, M.O., Whitwham, A., Keane, T., McCarthy, S.A., Davies, R.M., et al. (2021). Twelve years of SAMtools and BCFtools. GigaScience 10:giab008. 10.1093/gigascience/giab008.

Dinh, H.X., Singh, D., Periyannan, S., Park, R.F., and Pourkheirandish, M. (2020). Molecular genetics of leaf rust resistance in wheat and barley. Theoretical and Applied Genetics 133:2035–2050. 10.1007/s00122-020-03570-8.

Dinh, H.X., Singh, D., Gomez de la Cruz, D., Hensel, G., Kumlehn, J., Mascher, M., Stein, N., Perovic, D., Ayliffe, M., Moscou, M.J., et al. (2022). The barley leaf rust resistance gene Rph3 encodes a predicted membrane protein and is induced upon infection by avirulent pathotypes of Puccinia hordei. Nature Communications 13:2386. 10.1038/s41467-022-29840-1.

Dracatos, P.M., Singh, D., Bansal, U., and Park, R.F. (2015). Identification of new sources of adult plant resistance to Puccinia hordei in international barley (Hordeum vulgare L.) germplasm. European Journal of Plant Pathology 141:463–476. 10.1007/s10658-014-0556-9.

Dracatos, P.M., Barto¡, J., Elmansour, H., Singh, D., Karafiátová, M., Zhang, P., Steuernagel, B., Svačina, R., Cobbin, J.C.A., Clark, B., et al. (2019). The coiled-coil NLR Rph1, confers leaf rust resistance in barley cultivar Sudan Plant Physiology 179:1362–1372. 10.1104/pp.18.01052.

Elmansour, H., Singh, D., Dracatos, P.M., and Park, R.F. (2017). Identification and characterization of seedling and adult plant resistance to Puccinia hordei in selected African barley germplasm. Euphytica 213:119. 10.1007/s10681-017-1902-8.

Gilmour, J. (1973). Octal notation for designating physiologic races of plant pathogens. Nature 242:620–620. 10.1038/242620a0.

Golan, T., Anikster, Y., Moseman, J.G., and Wahl, I. (1978). A new virulent strain of Puccinia hordei. Euphytica 27:185–189. 10.1007/BF00039134.

Grabherr, M.G., Haas, B.J., Yassour, M., Levin, J.Z., Thompson, D.A., Amit, I., Adiconis, X., Fan, L., Raychowdhury, R., Zeng, Q., et al. (2011). Full-length transcriptome assembly from RNA-Seq data without a reference genome. Nature Biotechnology 29:644–652. 10.1038/nbt.1883.

Graner, A., Streng, S., Drescher, A., Jin, Y., Borovkova, I., and Steffenson, B.J. (2000). Molecular mapping of the leaf rust resistance gene Rph7 in barley. Plant Breeding 119:389–392. 10.1046/j.1439-0523.2000.00528.x.

Harwood, W.A. (2019). An introduction to barley: The crop and the model. Methods in molecular biology (Clifton, N.J.) 1900:1–5. 10.1007/978-1-4939-8944-7_1.

Holm, L. (2022). Dali server: structural unification of protein families. Nucleic acids research 50:W210–W215. 10.1093/nar/gkac387.

Jayakodi, M., Padmarasu, S., Haberer, G., Bonthala, V.S., Gundlach, H., Monat, C., Lux, T., Kamal, N., Lang, D., Himmelbach, A., et al. (2020). The barley pan-genome reveals the hidden legacy of mutation breeding. Nature 588:284–289. 10.1038/s41586-020-2947-8.

Jost, M., Singh, D., Lagudah, E., Park, R.F., and Dracatos, P. (2020). Fine mapping of leaf rust resistance gene Rph13 from wild barley. Theoretical and Applied Genetics 133:1887–1895. 10.1007/s00122-020-03564-6.

Jumper, J., Evans, R., Pritzel, A., Green, T., Figurnov, M., Ronneberger, O., Tunyasuvunakool, K., Bates, R., Žídek, A., Potapenko, A., et al. (2021). Highly accurate protein structure prediction with AlphaFold. Nature 596:583–589. 10.1038/s41586-021-03819-2.

Kim, D., Paggi, J.M., Park, C., Bennett, C., and Salzberg, S.L. (2019). Graph-based genome alignment and genotyping with HISAT2 and HISAT-genotype. Nature Biotechnology 37:907–915. 10.1038/s41587-019-0201-4.

Kim, D., Pertea, G., Trapnell, C., Pimentel, H., Kelley, R., and Salzberg, S.L. (2013). TopHat2: Accurate alignment of transcriptomes in the presence of insertions, deletions and gene fusions. Genome Biology 14:R36. 10.1186/gb-2013-14-4-r36.

Kolodziej, M.C., Singla, J., Sánchez-Martín, J., Zbinden, H., Šimková, H., Karafiátová, M., Doležel, J., Gronnier, J., Poretti, M., Glauser, G., et al. (2021). A membrane-bound ankyrin repeat protein confers race-specific leaf rust disease resistance in wheat. Nature Communications 12:956. 10.1038/s41467-020-20777-x.

Langmead, B., and Salzberg, S.L. (2012). Fast gapped-read alignment with Bowtie 2. Nature methods 9:357–359. 10.1038/nmeth.1923.

Li, B., and Dewey, C.N. (2011). RSEM: accurate transcript quantification from RNA-Seq data with or without a reference genome. BMC Bioinformatics 12:323. 10.1186/1471-2105-12-323.

Love, M.I., Huber, W., and Anders, S. (2014). Moderated estimation of fold change and dispersion for RNA-seq data with DESeq2. Genome Biology 15:550. 10.1186/s13059-014-0550-8.

Mascher, M., Wicker, T., Jenkins, J., Plott, C., Lux, T., Koh, C.S., Ens, J., Gundlach, H., Boston, L.B., Tulpová, Z., et al. (2021). Long-read sequence assembly: a technical evaluation in barley. The Plant cell 33:1888–1906. 10.1093/plcell/koab077.

Mehnaz, M., Dracatos, P.M., Dinh, H.X., Forrest, K., Rouse, M.N., Park, R.F., and Singh, D. (2022). A novel locus conferring resistance to Puccinia hordei maps to the genomic region corresponding to Rph14 on barley chromosome 2HS. Frontiers in Plant Science 13:980870. 10.3389/fpls.2022.980870.

Mehnaz, M., Dracatos, P., Pham, A., March, T., Maurer, A., Pillen, K., Forrest, K., Kulkarni, T., Pourkheirandish, M., Park, R.F., et al. (2021). Discovery and fine mapping of Rph28: a new gene conferring resistance to Puccinia hordei from wild barley. Theoretical and Applied Genetics 134:2167–2179. 10.1007/s00122-021-03814-1.

Monat, C., Padmarasu, S., Lux, T., Wicker, T., Gundlach, H., Himmelbach, A., Ens, J., Li, C., Muehlbauer, G.J., Schulman, A.H., et al. (2019). TRITEX: chromosome-scale sequence assembly of Triticeae genomes with open-source tools. Genome Biology 20:284. 10.1186/s13059-019-1899-5.

Park, R.F., and Karakousis, A. (2002). Characterization and mapping of gene Rph19 conferring resistance to Puccinia hordei in the cultivar ‘Reka 1’ and several Australian barleys. Plant Breeding 121:232–236. 10.1046/j.1439-0523.2002.00717.x.

Park, R.F., Golegaonkar, P.G., Derevnina, L., Sandhu, K.S., Karaoglu, H., Elmansour, H.M., Dracatos, P.M., and Singh, D. (2015). Leaf rust of cultivated barley: Pathology and control. Annual Review of Phytopathology 53:565–589. 10.1146/annurev-phyto-080614-120324.

Pettersen, E.F., Goddard, T.D., Huang, C.C., Meng, E.C., Couch, G.S., Croll, T.I., Morris, J.H., and Ferrin, T.E. (2021). UCSF ChimeraX: Structure visualization for researchers, educators, and developers. Protein Science 30:70–82. 10.1002/pro.3943.

Qi, X., Niks, R.E., Stam, P., and Lindhout, P. (1998). Identification of QTLs for partial resistance to leaf rust (Puccinia hordei) in barley. Theoretical and Applied Genetics 96:1205–1215. 10.1007/s001220050858.

Roane, C., and Starling, T. (1970). Inheritance of reaction to Puccinia hordei in barley. III. Genes in the cultivars Cebada Capa and Franger. Phytopathology 57:66–68. 10.1094/Phyto-60-788.

Rothwell, C.T., Singh, D., Dracatos, P.M., and Park, R.F. (2020). Inheritance and dharacterization of Rph27: A third race-specific resistance gene in the barley cultivar Quinn. Phytopathology 110:1067–1073. 10.1094/phyto-12-19-0470-r.

Sánchez-Martín, J., and Keller, B. (2021). NLR immune receptors and diverse types of non-NLR proteins control race-specific resistance in Triticeae. Current Opinion in Plant Biology 62:102053. 10.1016/j.pbi.2021.102053.

Sánchez-Martín, J., Steuernagel, B., Ghosh, S., Herren, G., Hurni, S., Adamski, N., Vrána, J., Kubaláková, M., Krattinger, S.G., Wicker, T., et al. (2016). Rapid gene isolation in barley and wheat by mutant chromosome sequencing. Genome Biology 17:221. 10.1186/s13059-016-1082-1.

Scherrer, B., Isidore, E., Klein, P., Kim, J.S., Bellec, A., Chalhoub, B., Keller, B., and Feuillet, C. (2005). Large intraspecific haplotype variability at the Rph7 locus results from rapid and recent divergence in the barley genome. Plant Cell 17:361–374. 10.1105/tpc.104.028225.

Shtaya, M.J.Y., Sillero, J.C., and Rubiales, D. (2006). Identification of a new pathotype of Puccinia hordei with virulence for the resistance gene Rph7. European Journal of Plant Pathology 116:103–106. 10.1007/s10658-006-9043-2.

Singh, D., Dracatos, P.M., Loughman, R., and Park, R.F. (2016). Genetic mapping of resistance to Puccinia hordei in three barley doubled-haploid populations. Euphytica 213:16. 10.1007/s10681-016-1799-7.

Singh, D., Ziems, L.A., Dracatos, P.M., Pourkheirandish, M., Tshewang, S., Czembor, P., German, S., Fowler, R.A., Snyman, L., Platz, G.J., et al. (2018). Genome-wide association studies provide insights on genetic architecture of resistance to leaf rust in a worldwide barley collection. Molecular Breeding 38:43. 10.1007/s11032-018-0803-4.

Steffenson, B.J.J. Y.; Griffey, C. A. (1993). Pathotypes of Puccinia hordei with virulence for the barley leaf rustresistance gene Rph7 in the United-States. Plant Disease 77:867–869. 10.1094/PD-77-0867.

Tuleen, N., and McDaniel, M. (1971). Location of genes Pa and Pa5. Barley Newsl 15:106–107.

Walkowiak, S., Gao, L., Monat, C., Haberer, G., Kassa, M.T., Brinton, J., Ramirez-Gonzalez, R.H., Kolodziej, M.C., Delorean, E., Thambugala, D., et al. (2020). Multiple wheat genomes reveal global variation in modern breeding. Nature 588:277–283. 10.1038/s41586-020-2961-x.

Wang, Y., Subedi, S., de Vries, H., Doornenbal, P., Vels, A., Hensel, G., Kumlehn, J., Johnston, P.A., Qi, X., Blilou, I., et al. (2019). Orthologous receptor kinases quantitatively affect the host status of barley to leaf rust fungi. Nature Plants 5:1129–1135. 10.1038/s41477-019-0545-2.

Welner, D.H., Lindemose, S., Grossmann, J.G., Møllegaard, N.E., Olsen, A.N., Helgstrand, C., Skriver, K., and Lo Leggio, L. (2012). DNA binding by the plant-specific NAC transcription factors in crystal and solution: a firm link to WRKY and GCM transcription factors. Biochemical Journal 444:395–404. 10.1042/bj20111742.

Zhang, J., Zhang, P., Dodds, P., and Lagudah, E. (2020). How target-sequence enrichment and sequencing (TEnSeq) pipelines have catalyzed resistance gene cloning in the wheat-eust pathosystem. Frontiers in Plant Science 11 10.3389/fpls.2020.00678.

Zhou, J.-M., and Zhang, Y. (2020). Plant immunity: Danger perception and signaling. Cell 181:978–989. 10.1016/j.cell.2020.04.028.

