## Supplementary File for "A pathogen-induced putative NAC transcription factor mediates leaf rust resistance in barley"

### Supplemental Figure 1

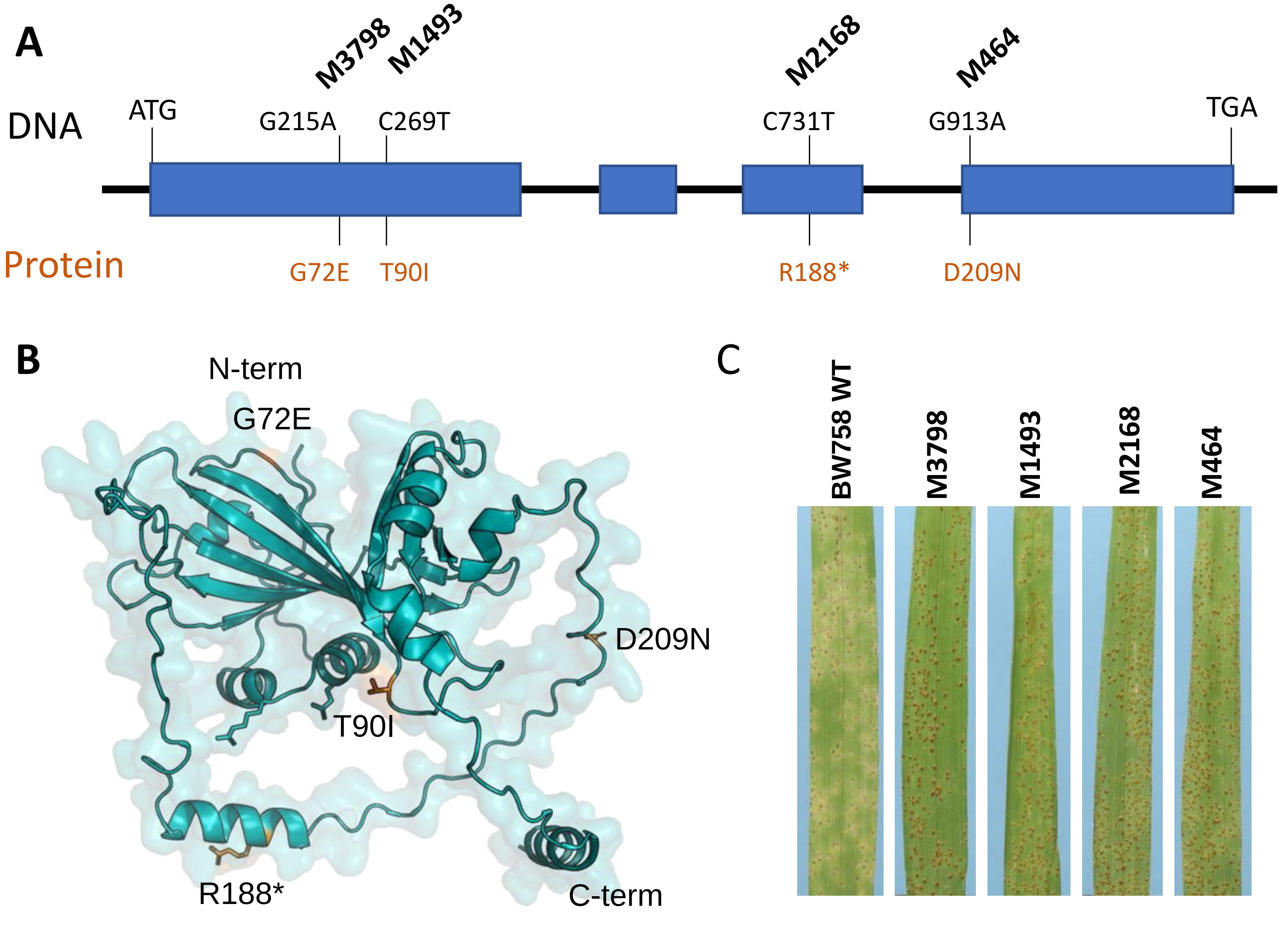

### Supplemental Figure 2

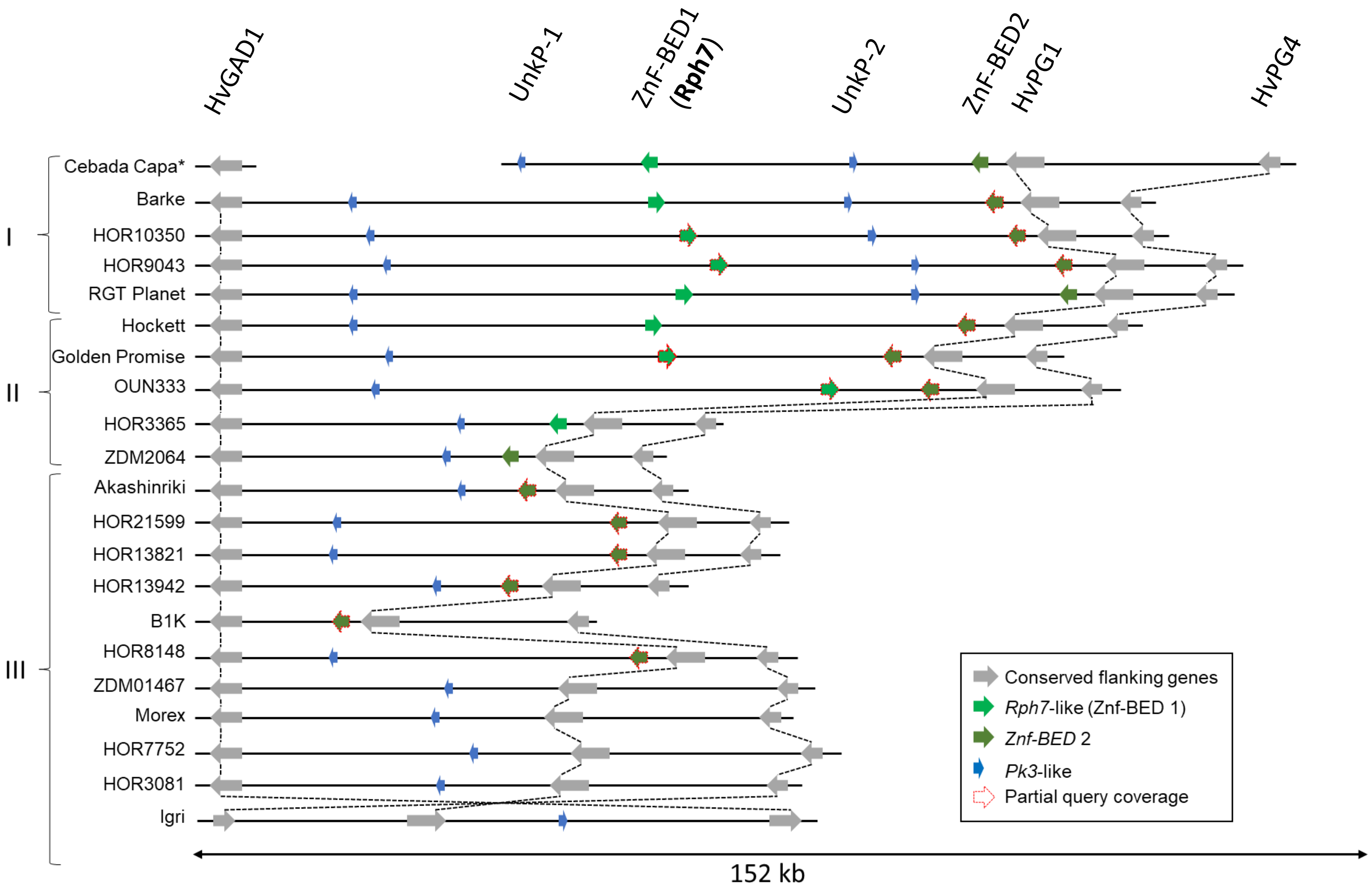

### Supplemental Figure 3

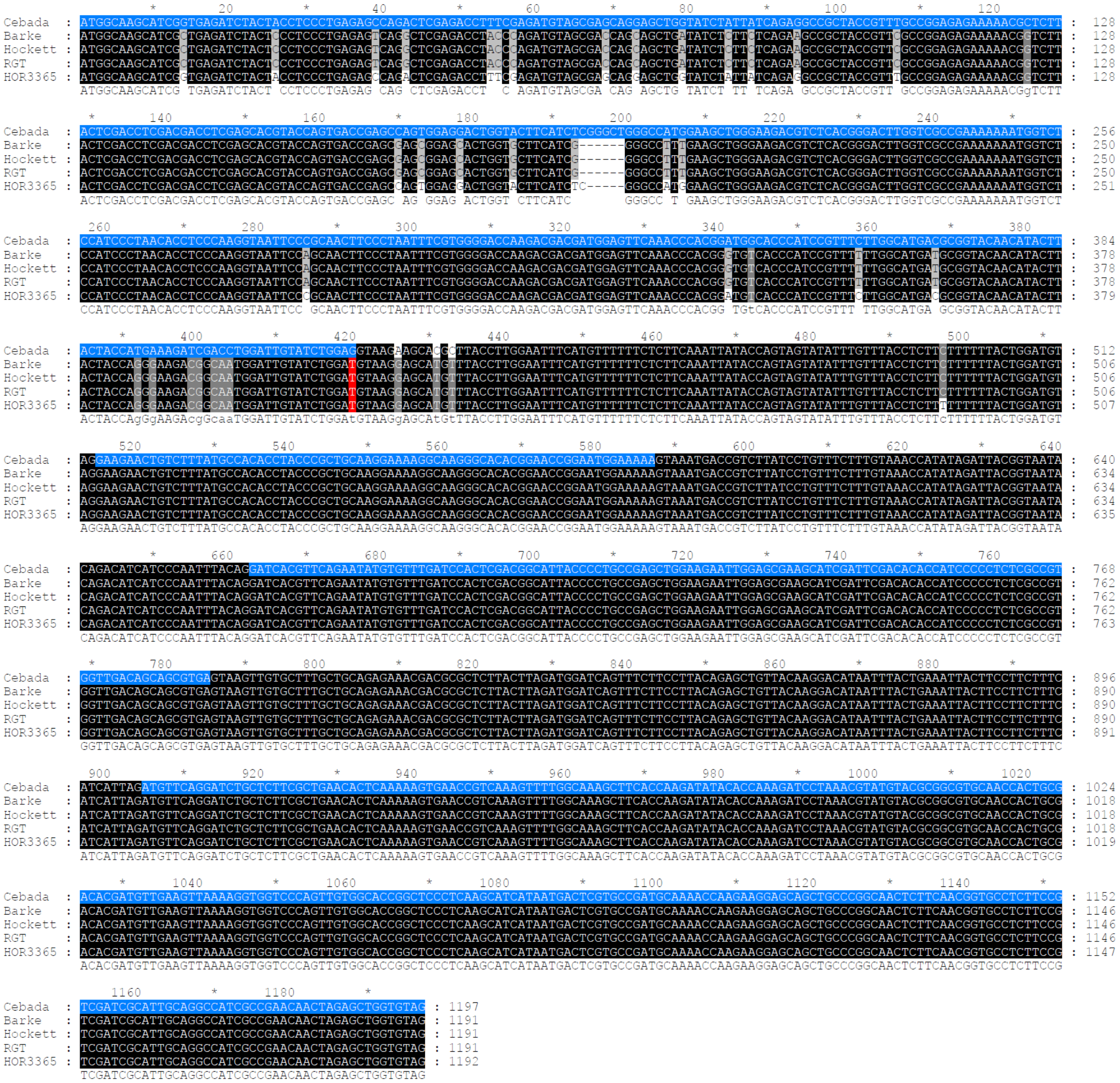

### Supplemental Figure 4

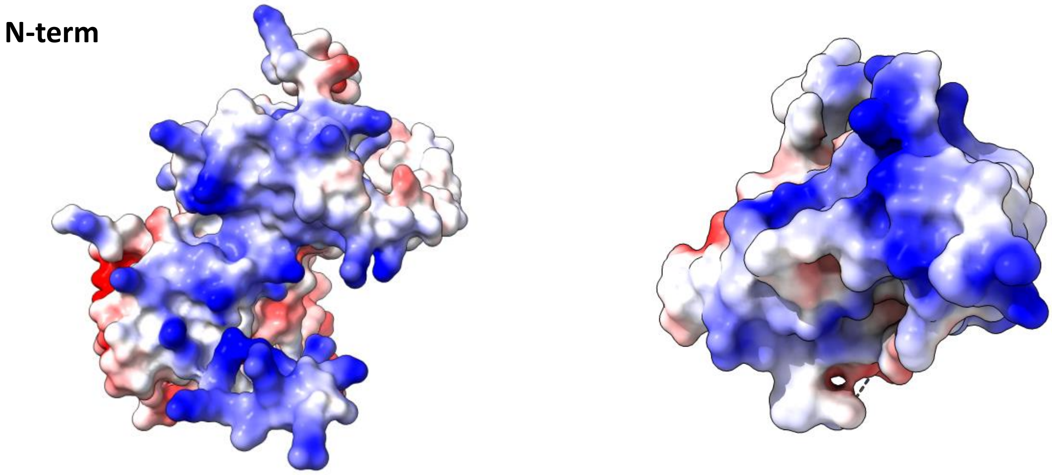

### Supplemental Figure 5

A

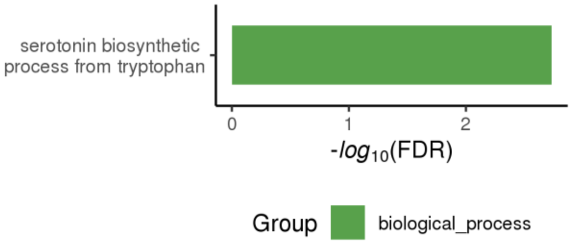


B

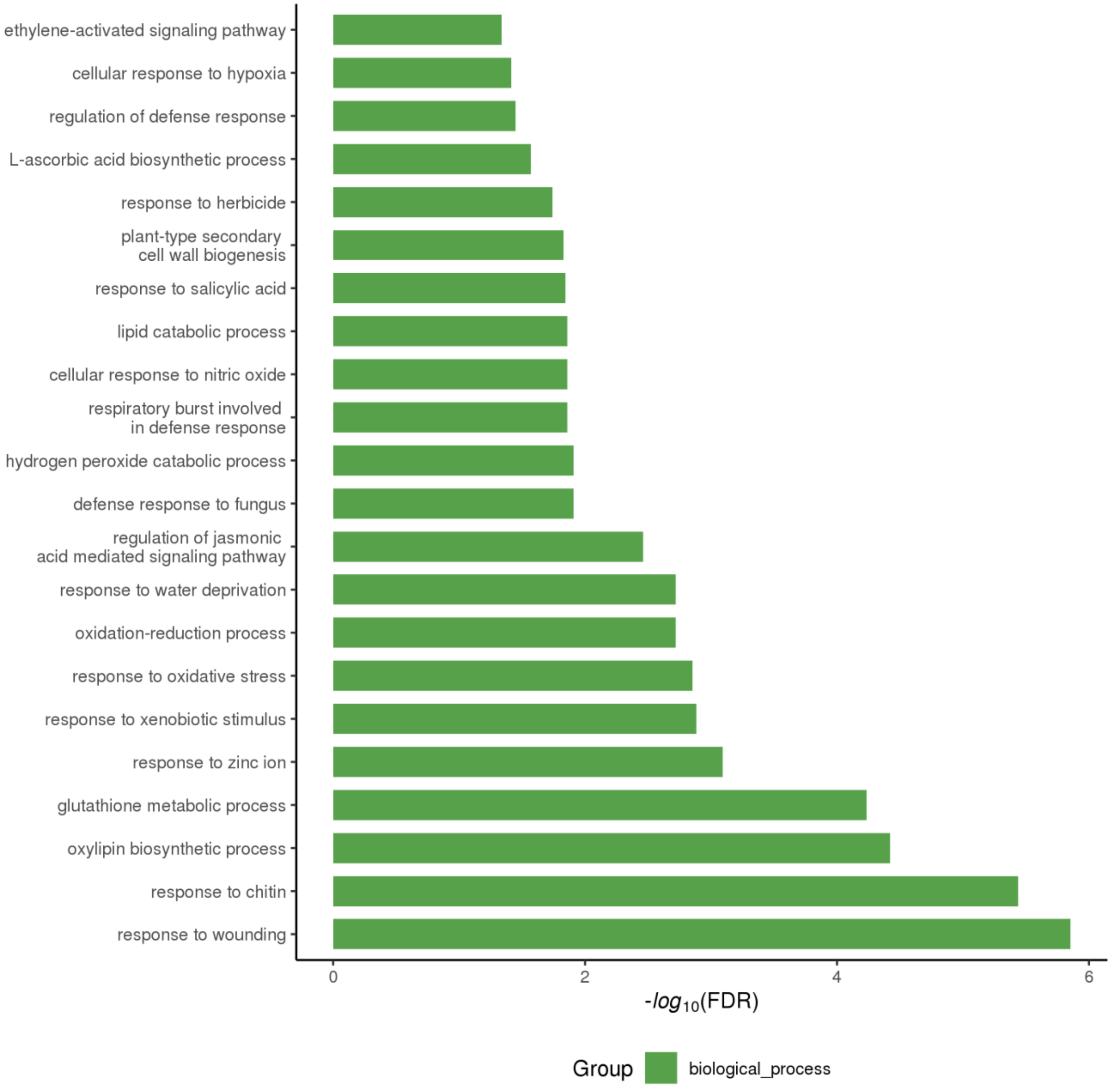

### Supplemental Figure 6

| Gene | log2FoldChange | Description | Group |
|---|---|---|---|
| HORVU.MOREX.r3.1HG0000480 | 11.5167168 | Jasmonate-induced protein | Jasmonate-related |
| HORVU.MOREX.r3.2HG0205090 | 8.957075061 | Jasmonate-induced protein | |
| HORVU.MOREX.r3.1HG0000390 | 8.798539053 | Jasmonate-induced protein | |
| HORVU.MOREX.r3.3HG0330760 | 8.368007039 | Jasmonate-induced protein | |
| HORVU.MOREX.r3.3HG0330600 | 7.945613979 | Jasmonate-induced protein | |
| HORVU.MOREX.r3.1HG0000450 | 7.481327173 | Jasmonate induced protein | |
| HORVU.MOREX.r3.3HG0330720 | 6.079099777 | Jasmonate-induced protein | |
| HORVU.MOREX.r3.1HG0000470 | 5.3668004 | Jasmonate induced protein | |
| HORVU.MOREX.r3.3HG0330650 | 5.336017871 | Jasmonate induced protein | |
| HORVU.MOREX.r3.1HG0000430 | 5.03548161 | Jasmonate-induced protein | |
| HORVU.MOREX.r3.3HG0330620 | 4.561429708 | Jasmonate-induced protein | |
| HORVU.MOREX.r3.2HG0163880 | 2.744873093 | Jasmonate zim-domain protein | |
| HORVU.MOREX.r3.4HG0405230 | 3.641368552 | Jasmonate ZIM domain protein | |
| HORVU.MOREX.r3.5HG0480600 | 2.450071629 | Jasmonate zim-domain protein | |
| HORVU.MOREX.r3.7HG0669350 | -2.248440698 | Jasmonate zim-domain protein | |
| HORVU.MOREX.r3.7HG0669360 | -3.843135333 | Jasmonate zim-domain protein | |
| HORVU.MOREX.r3.5HG0519340 | 7.849094936 | Pathogenesis-related protein 1 | Pathogenesis-related |
| HORVU.MOREX.r3.6HG0619530 | 3.726234428 | Pathogenesis-related protein PR-4 | |
| HORVU.MOREX.r3.7HG0662370 | 2.811237119 | Pathogenesis-related protein 1 | |
| HORVU.MOREX.r3.4HG0383530 | 2.535321426 | Pathogenesis-related protein 1 | |
| HORVU.MOREX.r3.5HG0423030 | 2.687585048 | Thaumatin-like protein | |
| HORVU.MOREX.r3.5HG0423110 | 2.23607062 | Thaumatin-like protein | |
| HORVU.MOREX.r3.5HG0423060 | 2.098756469 | Thaumatin-like protein | |
| HORVU.MOREX.r3.3HG0307860 | 9.475069647 | Salicylate O-methyltransferase | |
| HORVU.MOREX.r3.3HG0297450 | 5.927803876 | WRKY transcription factor | WRKY transcription factors |
| HORVU.MOREX.r3.5HG0464240 | 5.622365016 | WRKY transcription factor | |
| HORVU.MOREX.r3.3HG0237360 | 5.428512893 | WRKY family transcription factor | |
| HORVU.MOREX.r3.1HG0070910 | 4.086371185 | WRKY transcription factor | |
| HORVU.MOREX.r3.6HG0618760 | 3.920601689 | WRKY transcription factor, putative | |
| HORVU.MOREX.r3.7HG0710560 | 2.86203917 | WRKY transcription factor | |
| HORVU.MOREX.r3.1HG0088760 | 2.84158887 | WRKY transcription factor | |
| HORVU.MOREX.r3.1HG0092420 | 2.575085338 | WRKY transcription factor | |
| HORVU.MOREX.r3.1HG0070890 | 2.430739479 | WRKY family transcription factor | |
| HORVU.MOREX.r3.3HG0286660 | 2.227980799 | WRKY transcription factor | |
| HORVU.MOREX.r3.3HG0273660 | 2.08655692 | WRKY transcription factor | |
| HORVU.MOREX.r3.7HG0743280 | -2.521453205 | WRKY transcription factor | |
| HORVU.MOREX.r3.7HG0743270 | -2.649562839 | WRKY transcription factor | |
| HORVU.MOREX.r3.1HG0080940 | -2.717058589 | WRKY transcription factor | |
| HORVU.MOREX.r3.4HG0333220 | -5.75234814 | WRKY transcription factor | |
| HORVU.MOREX.r3.3HG0222490 | 5.583724968 | Disease resistance protein RPM1 | Disease resistance proteins |
| HORVU.MOREX.r3.2HG0214310 | 5.499532862 | Disease resistance protein RPM1 | |
| HORVU.MOREX.r3.3HG0316460 | 4.21074174 | Disease resistance protein (NBS-LRR class) family | |
| HORVU.MOREX.r3.2HG0106240 | 4.030136776 | disease resistance protein (TIR-NBS-LRR class) | |
| HORVU.MOREX.r3.5HG0514590 | 3.341479475 | NBS-LRR disease resistance protein, putative | |
| HORVU.MOREX.r3.6HG0611400 | 2.872707076 | NB-ARC domain-containing disease resistance protein | |
| HORVU.MOREX.r3.6HG0630620 | 2.853213892 | Disease resistance protein RGA2 | |
| HORVU.MOREX.r3.7HG0744510 | 2.29995067 | Disease resistance protein (NBS-LRR class) family | |
| HORVU.MOREX.r3.6HG0559440 | -2.154378054 | Disease resistance protein RPP13 | |
| HORVU.MOREX.r3.7HG0712390 | -2.270475687 | NBS-LRR disease resistance protein | |
| HORVU.MOREX.r3.7HG0639110 | -2.630017639 | Disease resistance protein (NBS-LRR class) family | |
| HORVU.MOREX.r3.5HG0489840 | -3.103512187 | Disease resistance protein RPP13 | |
